# Virulence patterns of oat crown rust in Australia - season 2022

**DOI:** 10.1101/2023.09.24.558528

**Authors:** Eva C. Henningsen, David Lewis, Duong T. Nguyen, Jana Sperschneider, Shahryar F. Kianian, Eric Stone, Peter N. Dodds, Melania Figueroa

## Abstract

*Puccinia coronata* f. sp. *avenae* (*Pca*) is an important foliar pathogen of oat which causes crown rust disease. The virulence profile of 48 *Pca* isolates derived from different locations in Australia was characterised using a collection of oat lines often utilised in rust surveys in the USA and Australia. This analysis indicates that *Pca* populations in Eastern Australia are broadly virulent, in contrast to the population in Western Australia (WA). Several oat lines/*Pc* genes are effective against all rust samples collected from WA, suggesting they may provide useful resistance in this region if deployed in combination. We identified 19 lines from the USA oat differential set that display disease resistance to *Pca* in WA, some in agreement with previous rust survey reports. We adopted the 10-letter nomenclature system to define oat crown rust races in Australia and compare the frequency of those virulence traits to published data from the USA. Based on this nomenclature, 42 unique races were detected among the 48 isolates, reflecting the high diversity of virulence phenotypes for *Pca* in Australia. Nevertheless, the *Pca* population in the USA is substantially more broadly virulent than that of Australia. Close examination of resistance profiles for the oat differential set lines after infection with *Pca* supports hypotheses of allelism or redundancy among *Pc* genes or the presence of several resistance genes in some oat differential lines. These findings illustrate the need to deconvolute the oat differential set using molecular tools.

## Manuscript body

Oat crown rust disease is caused by the basidiomycete fungus *Puccinia coronata* f. sp. *avenae* (*Pca*) (Nazareno et al. 2018). This disease is prevalent in oat growing regions across Australia and worldwide, raising the profile of this pathogen as one of global importance. Oat is important to Australia’s economy as it is used for milling, grazing, and feed hay (Cowman et al. 2021). Australia has a leading position in the production of high-quality milling oats, and in the 2021-2022 cropping season produced 1.7 million tonnes of oat (Australian Bureau of Statistics 2021). Regions known for their production of milling oat include Western Australia (WA), the Eyre and York Peninsulas of South Australia (SA), Western and North-eastern Victoria (VIC), and the Riverina and Central New South Wales (NSW). Forage oats are widely grown in Central and South Queensland (QLD) and Northern NSW and to some extent in WA. Oat production in WA is heavily focused on grain for feed and hay. It is estimated that 48% of Australia’s hay export comes from WA alone (Troup 2017). Given the negative impact of crown rust on plant growth, yield, grain weight, and palatability, the natural populations of *Pca* must be closely monitored to detect changes in virulence traits. The recent increases in market demand for Australian oats have justified additional research and development investments to support the industry (Cowman et al. 2021).

The evolution of virulence in *Pca* is driven by an arms race between the pathogen and the host (Nazareno et al. 2018). The boom-and-bust cycle illustrates this process, as the pressure exerted by the extensive adoption of an oat cultivar with a disease resistance (*R*) gene leads to the selection of rare variants in the pathogen that can overcome that specific resistant trait. This results in a frequency increase of that virulence trait in the rust population (Figueroa et al. 2023). Consequently, planting of that oat cultivar may become less common, as it is no longer resistant to the pathogen. Oat resistance to *Pca* follows the gene-for-gene concept initially described by Flor (1971), which is now more broadly understood to involve recognition of the pathogen by the plant immune system (Dodds 2023; Pitsili et al. 2020). Plant *R* genes generally encode intracellular immune receptor proteins mostly belonging to the nucleotide binding leucine rich repeat receptor (NLR) class. These receptors recognise pathogen ‘effector’ proteins, known as virulence (Avr) factors, that are delivered into host cells during infection to suppress host basal defences and facilitate infection (Figueroa et al. 2021; Petre et al. 2014). This type of resistance is known as race-specific resistance, which provides the framework utilised by plant pathologists to nominate races (pathotypes) of a pathogen. These pathogen races represent specific virulence profiles on a set of selected cultivars that collectively are referred to as a host differential set. At the molecular level, it is the combination of Avr factors (effectors) that defines a race. Each of the differential cultivars preferably contains a single race-specific *R* gene; however, that is often difficult to achieve due to the time commitment and investment required to generate isogenic lines and develop molecular markers to ensure *R* gene presence. Oat differential sets are used extensively to categorise *Pca* races (pathotypes) but differ substantially in their composition in different regions (Carson 2011; Chong et al. 2000; Nazareno et al. 2018).

Through a network of plant pathologists and industry representatives, we received and processed samples for 48 *Pca* isolates derived from oat producing areas in Australia (WA:20, NSW:11, VIC:9, SA:4, QLD:4) to investigate the virulence landscape of this pathogen in the country and create a foundational resource for future studies (**Table 1**). Rust samples were received as infected foliar tissue material and recovered by inoculation onto the widely susceptible cultivars Swan or Marvelous. Single pustule isolates were collected and amplified using standard techniques to determine virulence profiles (Miller et al. 2020). Rust isolate stocks are kept at –80°C for long term storage. Presently, there is no universal oat differential set or nomenclature system to facilitate an international monitoring system. Thus, we adopted an oat differential set utilised for annual rust surveys in the USA and added several oat cultivars that are often included in rust surveys conducted by Australian Cereal Rust Control Programme (**Table S1**). Oat lines from the USA were imported from the USDA-ARS (St. Paul, MN, USA) to the CSIRO’s quarantine facility (Black Mountain Laboratories, Canberra, Australia). The remaining oat lines were accessed through the Australian Grains Genebank (AGG) or breeders. A single seed descent increase was undertaken for each oat line using standard growth conditions; briefly, plants were grown at 23°C for 16 hours light and 18°C for 8 hours dark and fertilized at stem elongation and anthesis with Osmocote® 19-9-10+2MgO+TE all-purpose slow release (Everris International B.V., Heerlen, Netherlands). To determine virulence profiles, *Pca* isolates were inoculated onto the differential lines and infection scores were recorded at 12 dpi with race assignments (pathotypes) made using the North American 16-letter code (Carson 2011; Chong et al. 2000; Nazareno et al. 2018). Infection scores were converted to a 0-9 numeric scale for statistical comparisons (Table S2) (Miller et al. 2018, 2020).

**Table 1.**
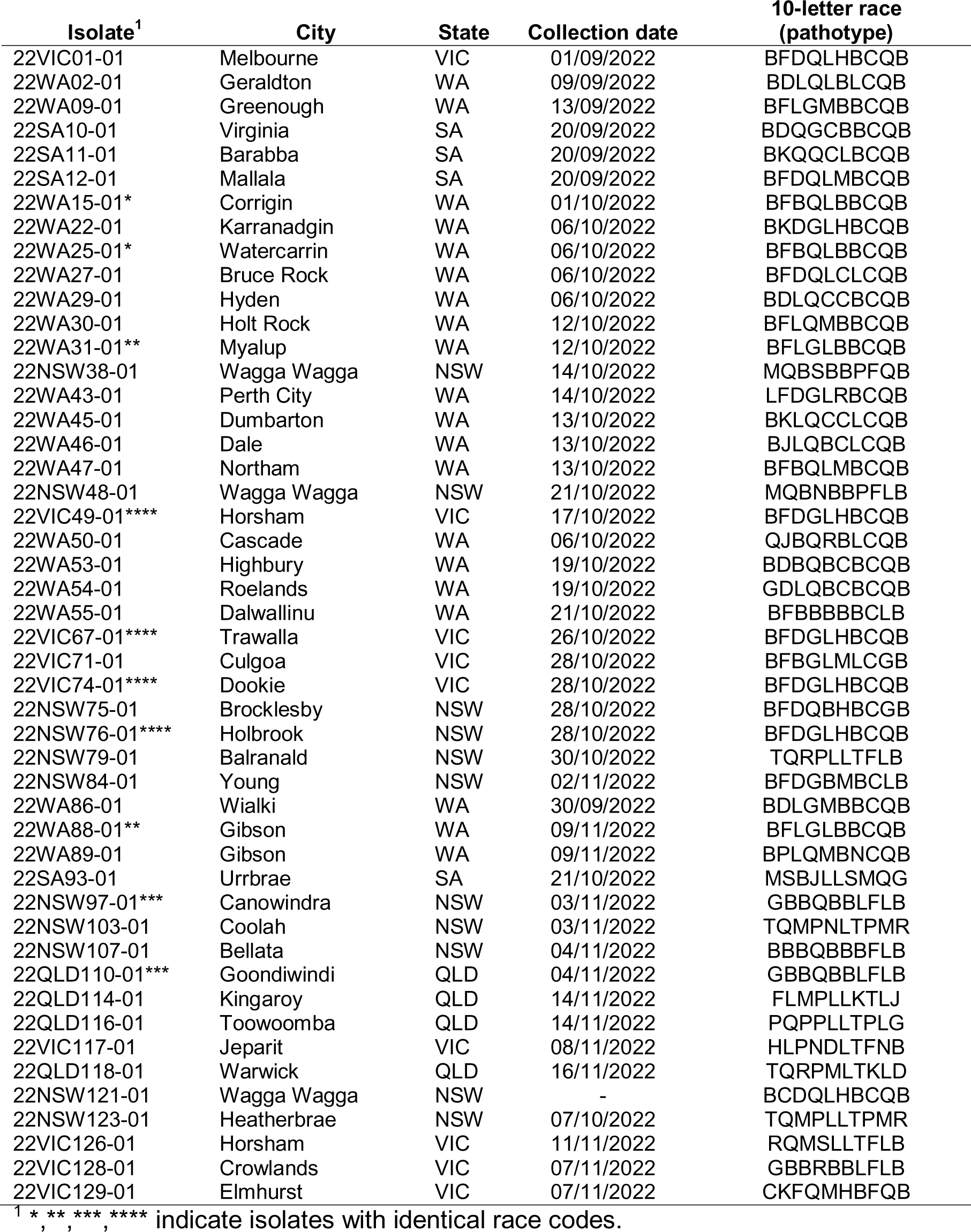
List of *Pca* isolates collected across Australia.

Overall, the *Pca* population from Eastern Australia (NSW, QLD, VIC, and SA) is more virulent compared to the population in Western Australia (**Figure 1A**; **Table S2**; Wilcoxon test p = 6.7 x 10^-8^). Collectively, *Pca* isolates from WA are only virulent on 21 of 40 USA differential lines, while isolates from Eastern Australia have virulence to 38 of 40 USA differential lines (**Figure 1B**). We identified 19 lines from the USA differential set that display resistance to all the *Pca* isolates collected from WA. Virulence to 18 lines, including Pc91, Pc94, and Pc96, was only sampled in Eastern Australia. Across the 31 USA differential lines with designated *Pc* genes, the most broadly virulent isolates were from eastern Australia (22QLD118, 22NSW103, 22NSW79), while the least broadly virulent isolates were from across Australia (22NSW107, 22WA53, 22WA86) (**Figure 1B**). Rust isolates from QLD had the highest average virulence on this subset, (5.50), followed by those from NSW (4.60), VIC (4.32), SA (4.29), with those from WA being least virulent on average (3.75) (**Table S2**). We obtained 42 unique 10-letter races for the entire *Pca* collection (48 isolates), illustrating the phenotypic diversity of the pathogen in Australia (**Table 1**).

**Figure 1.**
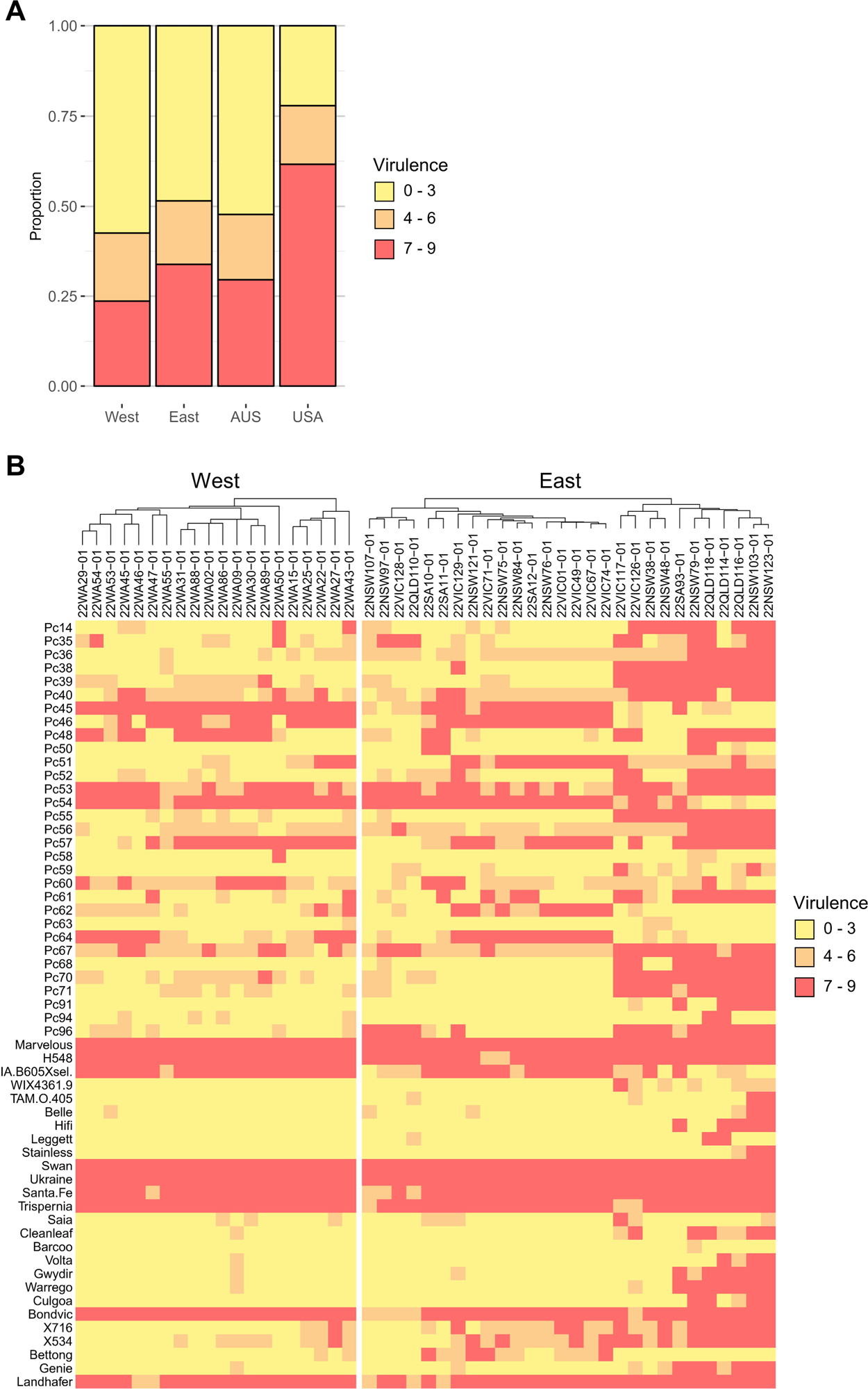
Virulence profile of a subset of 48 Australian *Puccinia coronata* f. sp. *avenae* (*Pca*) isolates from across the country. **A)** Barplots showing the proportion of cumulative infection types across the 40 USA differential lines for 48 Australian (West = WA; East = QLD, NSW, SA, and VIC) and 152 USA *Pca* isolates. **B)** Heatmap showing virulence profiles of isolates collected in 2022 (x-axis) on USA differential lines and cultivars used in Australian surveys (y-axis). High infection scores are shown in red indicating high virulence (susceptibility) and lower scores (yellow/orange) indicate avirulence (resistance). Columns are ordered by hierarchical clustering divided by region (West = WA; East = QLD, NSW, SA, and VIC). Heatmap was constructed using the R package ‘ComplexHeatmap’ (Gu 2016).

Virulence scores on 19 of the 40 USA differential lines can be compared with survey information from the Australian Cereal Rust Program (ACRCP) as they have been included in previous surveys (**Table S1**) (Australian Cereal Rust Survey 2020, 2021, 2022; Cuddy et al. 2016; Cuddy and Park 2014; Park 2013; Park and Kavanagh 2002, 2003, 2008, 2009, 2011; Park and Whale 1999). Of these, disease resistance scoring on 13 lines are consistent with reports from 1998-2022 by the ACRCP (Pc36, Pc39, Pc46, Pc50, Pc51, Pc56, Pc59, Pc61, Pc64, Pc68, Pc91, H548, WIX4361-9). Historically, virulence was detected in all 19 lines in Eastern Australia, but our study did not sample virulence for the Pc58 or Pc63 lines in this region (Cuddy et al. 2016; Cuddy and Park 2014; Park and Kavanagh 2009). We identified 12 lines which were resistant to all of the WA isolates in our sample, eight of which agree with historic survey records (Pc36, Pc50, Pc56, Pc59, Pc63, Pc68, Pc91, WIX4361-9), since virulence to these lines has never been recorded in WA. Virulence to the other four oat lines (Pc38, Pc52, Pc55, Pc71) has only been detected rarely in WA (Australian Cereal Rust Survey 2021, 2022; Park 2000), which may explain why no virulence was detected in this limited sample.

We also tested 17 oat lines often included as part of the ACRCP oat crown rust surveys (Brake et al. 2001; Park 2013). Six of these (Swan, Ukraine, Santa Fe, Trispernia, Bondvic, Landhafer) were susceptible to most *Pca* isolates from all Australian states. The *Pc* genes postulated to exist in these lines include *Pc3c*+*Pc4c*, *Pc4*, *Pc5*, *Pc6*, *Pc6c*, *Pc6d*, *Pc7*, *Pc8*, *Pc9*, and *Pc21*, which were deployed in the field more than 50 years ago (Nazareno et al. 2018; Simons 1985). As such, these lines are not viable sources of crown rust resistance and could be excluded from future crown rust surveys. Importantly, among the Australian lines, the cultivar Barcoo was the only genotype that was effective against all examined *Pca* isolates. However, according to historic records *Pca* overcame Barcoo’s resistance in 2001 (Park 2013), although this virulence trait seems to exist at a low frequency (only 1 or 2 samples each year) in NSW and QLD populations (Park 2013; Park and Wellings 2010; Park et al. 2022). There were eight other cultivars that displayed resistance to all WA isolates (Culgoa, Bettong, Cleanleaf, Saia, Volta, Warrego, Genie, Gwydir), consistent with survey data which detected no virulence to these lines in WA isolates (Australian Cereal Rust Survey 2020, 2021, 2022). The oat cultivar Cleanleaf is postulated to carry genes *Pc38*, *Pc39*, and *Pc52* (Park 2013), so our findings are consistent as we did not detect virulence on the Pc38 or Pc52 lines when testing *Pca* isolates from WA. However, the first report of crown rust virulence to Cleanleaf in eastern Australia (NSW and QLD) dates back to 1995 (Park et al. 2000). Similarly rust survey records indicate that virulence for cultivar Warrego was first identified in 1998, and in 2001 virulence on Gwydir and Bettong also emerged (Park 2013). Virulence on cultivars Genie and Volta are reported for years 2008 and 2010, respectively. These virulence traits are present in the pathogen populations of eastern Australia; therefore, it is possible for their frequency to increase or for virulent isolates to migrate to the west of the country.

Finally, we compared the phenotypic data for the Australian *Pca* isolates to data for 152 USA isolates collected between 2015-2018 (Hewitt et al. 2023; Miller et al. 2020) to compare with Australian data. As illustrated by the violin plots (**Figure 1A**), the 2022 Australian *Pca* collection is significantly less virulent than the USA collection from 2015-2018 (mean virulence 3 and 7.5, respectively; Wilcoxon test p < 2.2 x 10^-16^). By comparing virulence profiles between Australian *Pca* in this study and *Pca* from the USA (Hewitt et al. 2023; Miller et al. 2020), we noted that certain differential lines have already lost resistance in the USA but were still mostly effective in Australia (WIX4361-9, Belle, Stainless). Furthermore, the oat differential lines named Pc94 and Leggett, which is postulated to carry Pc68 and Pc94, still exhibit broad resistance in both countries at present, although we expect virulence evolution is already occurring in both the USA and Australia due to evolutionary pressure exerted by the use of these *R* genes.

Two GWAS analyses of virulence in *Pca* have indicated complex relationships among *Pc* genes represented in the oat differential sets (Hewitt et al. 2023; Miller et al. 2020). Noticeably, we observed highly similar resistance profiles for Pc39, Pc38, Pc55, Pc70, and Pc71 in our infection assays. Previously, the *R* genes *Pc*39, *Pc*55, and *Pc*71 have been postulated as being either allelic or the same gene by various authors (Chong and Seaman 1989; Kiehn et al. 1976; Leonard et al. 2005). Moreover, GWAS results (Hewitt et al. 2023; Miller et al. 2020) for USA *Pca* isolates detected virulence associations for Pc38, Pc39, Pc55, Pc63, Pc70, and Pc71 in the same genomic interval, suggesting that some of the immunoreceptors encoded by these genes recognise the same *Avr* effector or genetically linked *Avr* effectors. Thus, data derived from the Australian *Pca* infections is consistent with this scenario. Furthermore, our results support the hypothesis by Hewitt et al. (2023) that the oat differential lines Pc62 and Pc64 carry alleles of the same *R* gene as the resistance profiles of these lines have overlap suggesting allelism as observed in the USA *Pca* population. The resistance phenotypes of the lines Pc35 and Pc58 also agree with the results from Hewitt et al. (2023), suggesting that the oat lines Pc58 and Pc35 carry one *R* gene in common and the Pc58 line contains at least one additional *R* gene. As highlighted previously by Hewitt et al. (2023), we also observed consistency between the resistance profiles of the oat differential line Pc91 and the oat cultivar Hi-Fi, which was derived from Amagalon, the original source of *Pc91* (McMullen et al. 2005). In summary, our findings provide additional motivation for the need to develop an oat differential set that clearly differentiates among *R* genes and eliminates redundancy.

The expansion of the oat crown rust collection in the coming years coupled with population genomics analysis will provide an accurate representation of the pathogen’s genetic diversity. Recent advances in generating genome references of *Pca* (Miller at al. 2018) including a nuclear-phased chromosome-level assembly (Henningsen et al. 2022) provide a strong foundation to study the genetic relationships of these isolates. Such information will be instrumental in determining the factors contributing to host adaptation of *Pca* in Australia. Rust fungi are known to evolve virulence by mutation, reassortment of virulence alleles, and somatic hybridisation (Figueroa et al. 2020). The invasive species *Rhamnus cathartica*, also known as common buckthorn, acts as an alternate host for *Pca* and allows sexual reproduction (Nazareno et al. 2018). Several studies (Berlin et al. 2018; Hewitt et al. 2023; Miller et al. 2020; Zhao et al. 2016) document the influence of sexuality in the diversity of the oat crown rust pathogen. In Australia, a sexual host for *Pca* has not been reported (Burdon and Thrall 2008), so sexual reproduction is not expected to play a role in the evolution of *Pca* in Australia, suggesting the Australian population is evolving by clonality and stepwise mutation. Wild oats are abundant in the Australian landscape and serve as an additional host for *Pca*, which is predicted to favour mutations and contribute to the emergence of diversity (Burdon and Thrall 2008). However, it is not possible to define lineages from the phenotypic data alone; more detailed genotypic analysis will be required to fully characterize the number and diversity of clonal lineages present in the Australian *Pca* population.

The findings from this study, along with recent research (Hewitt et al. 2023; Miller et al. 2020) highlight the importance in developing durable crown rust resistance in oat. This can be achieved by stacking the most current effective genes, including both APR (Adult Plant Resistance) and ASR (All Stages Resistance) (Periyannan et al. 2017) and identifying and integrating novel sources of resistance into the oat breeding pools (Figueroa et al. 2020; Klos et al. 2017; Nazareno et al. 2018, 2023).

## Acknowledgments

We thank Ben Trevaskis and Meredith McNeil at CSIRO, as well as Bruce Winter at the Department of Agriculture and Fisheries (Queensland) for supplying seed for the oat cultivars Cleanleaf, Gwydir, Warrego, Barcoo, and Genie. We would also like to thank Liza Apps for technical support at the CSIRO’s Quarantine facility. Finally, we also thank Allan Rattey (InterGrain), Mark McLean (Agriculture Victoria), Hari Dadu (Agriculture Victoria), Ciara Beard (DPIRD WA^1^), Kylie Chambers (DPIRD WA), Andrea Hills (DPIRD WA), Jason Bradley (DPIRD WA), Joel Kidd (DPIRD WA), Tara Garrard (SARDI^2^), Brad Baxter (NSW DPI^3^), Mia Bowen Osmond (Palafor Partners Pty. Ltd.), Lee Hickey (University of Queensland), and Peter Dracatos (La Trobe University) and their organizations for contributing samples to the collection used in this study.

## Footnotes

1 Department of Primary Industries and Regional Development, WA

2 South Australian Research and Development Institute

3 New South Wales Department of Primary Industries

## Literature Cited

Agricultural Commodities, Australia. 2023. Australian Bureau of Statistics. https://www.abs.gov.au/statistics/industry/agriculture/agricultural-commodities-australia/latest-release

Australian Cereal Rust Survey. 2020. Plant Breeding Institute, University of Sydney. https://www.google.com/maps/d/viewer?mid=1VZPy5uGhC9RXfgp4TUuwHEm0H8QXRSWJ&ll=-32.93953349389678%2C122.88331689066194&z=7

Australian Cereal Rust Survey. 2021. Plant Breeding Institute, University of Sydney. https://www.google.com/maps/d/viewer?mid=17k2hAS9ProHR8c9DiAPlWJEUeoys5WLM&ll=-30.936465508880048%2C121.47685631082572&z=7

Australian Cereal Rust Survey. 2022. Plant Breeding Institute, University of Sydney. https://www.google.com/maps/d/viewer?mid=1qzwnH1u0B2apVvpkEKFm0mfBP_7cBknF&ll=-28.28748075112555%2C133.1960683213396&z=6

Berlin, A., Wallenhammar, A. C., and Andersson, B. 2018. Population differentiation of *Puccinia coronata* between hosts –implications for the epidemiology of oat crown rust. Eur. J. Plant Pathol. 152:901–907.

Brake, V. M., Irwin, J. A. G., and Park, R. F. 2001. Genetic variability in Australian isolates of *Puccinia coronata* f. sp. *avenae* assessed with molecular and pathogenicity markers. Australas. Plant Pathol. 30:259–266.

Burdon, J. J., and Thrall, P. H. 2008. Pathogen evolution across the agro-ecological interface: implications for disease management. Evol. Appl. 1:57–65.

Carson, M. L. 2011. Virulence in oat crown rust (*Puccinia coronata* f. sp. *avenae*) in the United States from 2006 through 2009. Plant Dis. 95:1528–1534.

Chong, J., Leonard, K. J., and Salmeron, J. J. 2000. A North American system of nomenclature for *Puccinia coronata* f. sp. *avenae*. Plant Dis. 84:580–585.

Chong, J., and Seaman, W. L. 1989. Virulence and distribution of *Puccinia coronata* in Canada in 1988. Can. J. Plant Pathol. 11:439–442.

Chong, J., and Zegeye, T. 2004. Physiologic specialization of *Puccinia coronata* f. sp. *avenae*, the cause of oat crown rust, in Canada from 1999 to 2001. Can. J. Plant Pathol. 26:97–108.

Cowman, S., Cox, B., Yamamoto, M., and Kingwell, R. 2021. Opportunities and risks for the Australian oats industry. Australian Export Grains Innovation Centre.

Cuddy, W., and Park, R. F. 2014. Cereal rust situation update, October 2014. Cereal Rust Report 12(4). Plant Breeding Institute, University of Sydney.

Cuddy, W., Park, R. F., and Singh, D. 2016. Cereal rust situation, September 2016. Cereal Rust Report 14(7). Plant Breeding Institute, University of Sydney.

Dodds, P. N. 2023. From gene-for-gene to resistosomes: Flor’s enduring legacy. MPMI. 36(8):461–467.

Figueroa, M., Dodds, P. N., and Henningsen, E. C. 2020. Evolution of virulence in rust fungi — multiple solutions to one problem. Curr. Opin. Plant Biol. 56:20–27.

Figueroa, M., Dodds, P. N., Henningsen, E. C., and Sperschneider, J. 2023. Global landscape of rust epidemics by *Puccinia* species: Current and future perspectives. In Plant Relationships: Fungal-Plant Interactions, eds. Barry Scott and Carl Mesarich. Cham: Springer International Publishing, p. 391–423.

Figueroa, M., Ortiz, D., and Henningsen, E. C. 2021. Tactics of host manipulation by intracellular effectors from plant pathogenic fungi. Curr. Opin. Plant Biol. 62:102054

Flor, H. H. 1971. Current status of the gene-for-gene concept. Annu. Rev. Phytopathol. 9:275–296.

Gu, Z., Eils, R., and Schlesner, M. 2016. Complex heatmaps reveal patterns and correlations in multidimensional genomic data. Bioinformatics. 32:2847–2849.

Henningsen, E. C., Hewitt, T., Dugyala, S., Nazareno, E. S., Gilbert, E., Li, F., Kianian, S. F., Steffenson, B. J., Dodds, P. N., Sperschneider, J., and Figueroa, M. 2022. A chromosome-level, fully phased genome assembly of the oat crown rust fungus *Puccinia coronata* f. sp. *avenae*: a resource to enable comparative genomics in the cereal rusts. G3. 12(8):jkac149.

Hewitt, T., Henningsen, E. C., Pereira, D., McElroy, K., Nazareno, E. S., Dugyala, S., Nguyen-Phuc, H., Li, F., Miller, M. E., Visser, B., Pretorius, Z., Boshoff, W., Sperschneider, J., Stukenbrock, E., Kianian, S. F., Dodds, P. N., and Figueroa, M. 2023. Genome-enabled analysis of population dynamics and virulence associated loci in the oat crown rust fungus *Puccinia coronata* f. sp. *avenae*. MPMI 10.1094/MPMI-09-23-0126-FI

Kiehn, F. A., McKenzie, R. I. H., and Harder, D. E. 1976. Inheritance of resistance to *Puccinia coronata avenae* and its association with seed characteristics in four accessions of *Avena sterilis*. Can. J. Genet. Cytol. 18(4):717–726

Klos, K. E., Yimer, B. A., Babiker, E. M., Beattie, A. D., Bonman, J. M., Carson, M. L., Chong, J., Harrison, S. A., Ibrahim, A. M. H., Kolb, F. L., McCartney, C. A., McMullen, M., Fetch, J. M., Mohammadi, M., Murphy, J. P., and Tinker, N. A. 2017. Genomelwide association mapping of crown rust resistance in oat elite germplasm. The Plant Genome. 10(2).

Leonard, K. J., Huerta-Espino, J., Salmeron, J. J. 2005. Virulence of oat crown rust in Mexico. Plant Dis. 89(9):941–948.

McMullen, M. S., Doehlert, D. C., and Miller, J. D. 2005. Registration of “HiFi” oat. Crop Sci. 45:1664–1665.

Miller, M. E., Nazareno, E. S., Rottschaefer, S. M., Riddle, J., Pereira, D. D. S., Li, F., Nguyen-Phuc, H., Henningsen, E. C., Persoons, A., Saunders, D. G. O., Stukenbrock, E., Dodds, P. N., Kianian, S. F., and Figueroa, M. 2020. Increased virulence of *Puccinia coronata* f. sp. *avenae* populations through allele frequency changes at multiple putative *Avr* loci. PLoS Genet. 16:e1009291.

Miller, M. E., Ying, Z., Vahid, O., Jana, S., Benjamin, S., Castle, R., Palmer, J. M., Garnica, D., Upadhyaya, N., Rathjen, J., Taylor, J. M., Park, R. F., Dodds, P. N., Hirsch, C. D., Kianian, S. F., and Figueroa, M. 2018. *De novo* assembly and phasing of dikaryotic genomes from two isolates of *Puccinia coronata* f. sp. *avenae*, the causal agent of oat crown rust. mBio. 9:10.1128/mbio.01650-17.

Nazareno, E. S., Fiedler, J. D., Ardayfio, N. K., Miller, M. E., Figueroa, M., and Kianian, S. F. 2023. Genetic analysis and physical mapping of oat adult plant resistance loci against *Puccinia coronata* f. sp. *avenae*. Phytopathology. 113:1307– 1316.

Nazareno, E. S., Li, F., Smith, M., Park, R. F., Kianian, S. F., and Figueroa, M. 2018. *Puccinia coronata* f. sp. *avenae*: a threat to global oat production. Mol. Plant Pathol. 19:1047–1060.

Park, R. F. 2013. New oat crown rust pathotype with virulence for *Pc91*. Cereal Rust Report 11(1). Plant Breeding Institute, University of Sydney.

Park, R. F. 2000. Occurrence and pathogenic specialisation in Puccinia coronata in Australasia, 1999-2000. Oat Newsletter. 46.

Park, R. F., Chhetri, M., Singh, D., and Ding, Y. 2022. Cereal rust situation, August 2022. Cereal Rust Report 19(2). Plant Breeding Institute, University of Sydney.

Park, R. F., and Kavanagh, P. 2002. 2001-2002 Cereal rust survey annual report: Oat leaf rust. Plant Breeding Institute, University of Sydney.

Park, R. F., and Kavanagh, P. 2003. 2002-2003 Cereal rust survey annual report: Oat leaf rust. Plant Breeding Institute, University of Sydney.

Park, R. F., and Kavanagh, P. 2008. 2007-2008 Cereal rust survey annual report: Oat crown rust. Plant Breeding Institute, University of Sydney.

Park, R. F., and Kavanagh, P. 2009. 2008-2009 Cereal rust survey annual report: Oat crown rust. Plant Breeding Institute, University of Sydney.

Park, R. F., and Kavanagh, P. 2011. 2010-2011 Cereal rust survey annual report: Oat crown rust. Plant Breeding Institute, University of Sydney.

Park, R. F., Oates, J. D., and Meldrum, S. 2000. Recent pathogenic changes in the leaf (brown) rust pathogen of wheat and the crown rust pathogen of oats in Australia in relation to host resistance. Acta Phytopathol. Entomol. Hung. 35:387–394.

Park, R. F. and Wellings, C. 2010. Cereal rust situation update, late spring 2010 2010. Cereal Rust Report 8(8). Plant Breeding Institute, University of Sydney.

Park, R. F., and Whale, M. 1999. 1998-1999 Cereal rust survey annual report: Oat leaf rust. Plant Breeding Institute, University of Sydney.

Periyannan, S., Milne, R. J., Figueroa, M., Lagudah, E. S., and Dodds, P. N. 2017. An overview of genetic rust resistance: From broad to specific mechanisms. PLoS Pathog. 13:e1006380.

Petre, B., Joly, D. L., and Duplessis, S. 2014. Effector proteins of rust fungi. Front. Plant Sci. 5:416.

Pitsili, E., Phukan, U. J., and Coll, N. S. 2020. Cell death in plant immunity. Cold Spring Harb. Perspect. Biol. 12:a036483.

Simons, M. D. 1985. Crown Rust. In: Diseases, Distribution, Epidemiology, and Control (pp. 131–172). Elsevier.

Troup, G. 2017. Hay exports. Department of Primary Industries and Regional Development, WA. https://www.agric.wa.gov.au/hay-production/hay-exports

Wickham, H. 2016. ggplot2: Elegant Graphics for Data Analysis. New York: Springer-Verlag.

Zhao, J., Wang, M., Chen, X., and Kang, Z. 2016. Role of alternate hosts in epidemiology and pathogen variation of cereal rusts. Annu. Rev. Phytopathol. 54:207–228.

